# Domestication of the emblematic white cheese-making fungus *Penicillium camemberti* and its diversification into two varieties

**DOI:** 10.1101/2020.02.12.945238

**Authors:** Jeanne Ropars, Estelle Didiot, Ricardo C. Rodríguez de la Vega, Bastien Bennetot, Monika Coton, Elisabeth Poirier, Emmanuel Coton, Alodie Snirc, Stéphanie Le Prieur, Tatiana Giraud

**Author notes:** **Corresponding author:** Jeanne Ropars.

## Abstract

Domestication involves recent adaptation under strong human selection and rapid diversification, and therefore constitutes a good model for studies of these processes. We studied the domestication of the emblematic white mold *Penicillium camemberti*, used for the maturation of soft cheeses, such as Camembert and Brie, about which surprisingly little was known, despite its economic and cultural importance. Whole genome-based analyses of genetic relationships and diversity revealed that an ancient domestication event led to the emergence of the gray-green *P. biforme* mold used in cheese-making, by divergence from the blue-green wild *P. fuscoglaucum* fungus. Another much more recent domestication event led to the generation of the *P. camemberti* clonal lineage as a sister group to *P. biforme. Penicillium biforme* displayed signs of phenotypic adaptation to cheese-making relative to *P. fuscoglaucum*, in terms of whiter color, faster growth on cheese medium under cave conditions, lower levels of toxin production and greater ability to prevent the growth of other fungi. The *P. camemberti* lineage displayed even stronger signs of domestication for all these phenotypic features. We also identified two differentiated *P. camemberti* varieties, apparently associated with different kinds of cheeses, and with contrasted phenotypic features in terms of color, growth, toxin production and competitive ability. We have, thus, identified footprints of domestication in these fungi, with genetic differentiation between cheese and wild populations, bottlenecks and specific phenotypic traits beneficial for cheese-making. This study has not only fundamental implications for our understanding of domestication but can also have important impacts on cheese-making.

## Introduction

Understanding how organisms adapt to their environment is a key issue in evolutionary biology, requiring investigations of population subdivision, and levels of genetic and phenotypic diversity or adaptive divergence. Domestication is a good model for studies of adaptive divergence, as it involves recent adaptation events affecting known traits under strong human selection and rapid diversification. Several studies on domesticated animals (e.g. horse, dog, pig [1–3]) and plants (e.g. maize, apricot [4,5]) have improved our understanding of adaptive divergence. Maize, for example, has undergone major changes in phenotype compared to its wild relative (teosinte), including a decrease in tillering, and the development of larger, non-dehiscent grains [6]. Different maize varieties have been selected for different usages, with sugar-rich varieties grown for human consumption as kernels and field corn varieties grown for animal feed. Such notable adaptation is often associated with a decrease in fitness in natural environments, with, for example, a decrease in or loss of sexual reproduction ability in bulldogs [7] and bananas [8].

Fungi are excellent models for studying evolution and adaptation in eukaryotes, given their many experimental assets [9], including their small genomes and tractability for laboratory experiments. Humans have domesticated several fungi for the fermentation of foods (e.g. for beer, bread, wine and cheese) [10]. Despite their economic importance, fungi used by humans have been little studied, with the exception of the budding yeast *Saccharomyces cerevisiae* used for beer, wine and bread production [11–21], the filamentous fungus *Aspergillus oryzae* used to ferment soy and rice products in Asia [22–24] and the blue-cheese mold *Penicillium roqueforti* [25–27]. Whole-genome analyses have revealed that *P. roqueforti* has been domesticated twice, resulting in one population specific to the Roquefort protected designation of origin (PDO), the other population being used worldwide for all types of blue cheeses [25–28]. The Roquefort population displays some genetic diversity and harbors beneficial traits for cheese production before the industrial era. By contrast, the non-Roquefort cheese population is a clonal lineage with traits beneficial for industrial cheese production, such as high levels of lipolytic activity, efficient cheese cavity colonization and high salt tolerance. Some of these beneficial traits have been conferred by two large horizontally transferred genomic regions, *Wallaby* and *CheesyTer* [26,27], in the non-Roquefort cheese population. A cheese-specific population also occurs in *S. cerevisiae*, differentiated from the populations used for alcohol or bread production, assimilating galactose more rapidly than *S. cerevisiae* populations thriving in other food environments (e.g. beer, bread, wine) or in natural environments (oak) [29].

The white mold *Penicillium camemberti* is used for the maturation of soft cheeses, such as Camembert, Brie and Neufchatel, and to a lesser extent for dry sausage maturation. It is thought to be a white mutant selected from the gray-green species *P. commune* for its color at the start of the 20th century [30], and cultured clonally ever since. The first records of Brie cheeses date from middle ages [31], being of blue aspect until the middle of the 20th century [32,33], as illustrated by a 19th century painting by Marie Jules Justin entitled “Symphonie des fromages en Brie majeur - Nature morte au fromage” (see graphical abstract) and a picture from the record “Fromages de France” published in 1953 [31]. Very little is known about the *P. camemberti* taxonomic status, origin and diversity, despite its great economic and cultural importance. In particular, its relationships to the closely related species *P. biforme, P. caseifulvum, P. commune* and *P. fuscoglaucum,* and even possible overlaps with these species, remain unclear. *Penicillium camemberti* has only ever been found in the food environment. Strains identified as *P. commune* are used for the maturation of other types of cheese (e.g. hard and blue cheeses) and in the production of dried sausages, and can be commonly found as spoilers of dairy products [34] and also in non-food environments. Genetic analyses however suggested that *P. commune* was not monophyletic (Figure 1A), which supported the reinstatement of two ancient species names, *P. biforme* and *P. fuscoglaucum* [35]. However, the most recent taxonomic reference study of the *Penicillium* genus maintained the *P. commune* name [36]. *Penicillium caseifulvum* has also sometimes been advocated to constitute a separate species in this clade [36], on the basis of colony morphology and the lack of production of cyclopiazonic acid (CPA), a mycotoxin known to be produced by *P. camemberti*. However, a study based on a few genetic markers was unable to differentiate these putative *P. caseifulvum* strains from *P. camemberti* [35].

**Figure 1:**
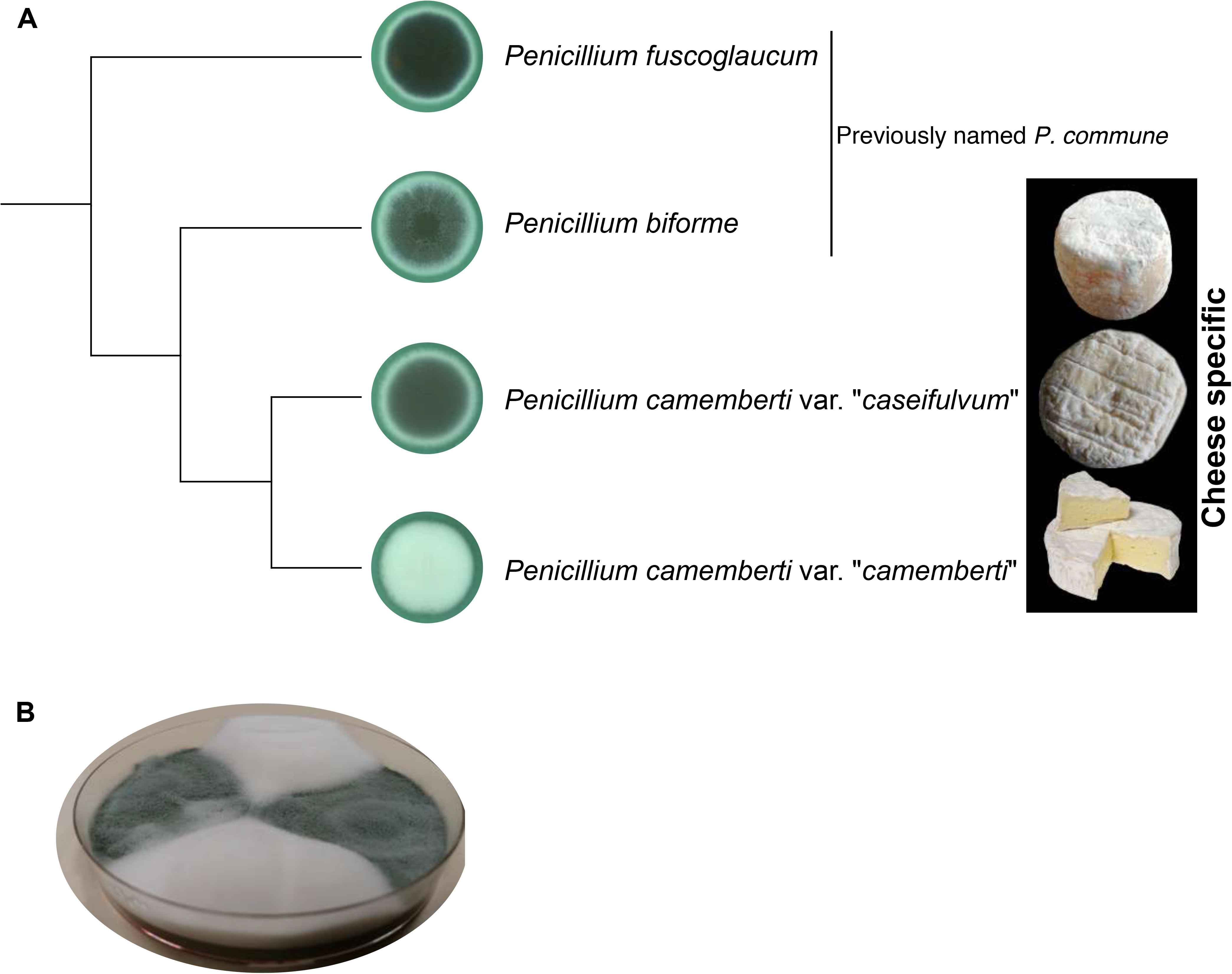
Cheeses and fungi used for cheese-making. A) Schematic representation of relationships between *Penicillium camemberti* and its closest relatives, *P. biforme* and *P. fuscoglaucum*. Pictures of *P. biforme, P. camemberti* var. “*camemberti*” and *P. camemberti* var. “*caseifulvum*” on salted cheese medium at 25°C after 10 days of growth. B) Petri dish with a white and fluffy *P. camemberti* var. “*camemberti*” strain (FM013) and a green rough *P. fuscoglaucum* strain (LCP06617) on malt medium at 25°C.

The taxonomy of this clade thus remains unclear, and population structure has not been studied with powerful genetic markers. Differences in phenotypic traits have not been extensively studied either. Humans may have selected strains for specific traits beneficial for cheese production, such as faster growth in cheese, attractive color and a greater ability to compete against food-spoiling microorganisms, but without the production of detrimental mycotoxins. Mycotoxins are natural secondary metabolites produced by fungi under certain conditions that can inhibit competitor growth and which can have a toxicological impact on humans [37]. Cyclopiazonic acid is one of these toxic secondary metabolites produced by *P. camemberti* under certain conditions [37,38].

Our goal here was to investigate the footprints of domestication by assessing genetic and phenotypic differentiation, by sequencing the genomes of strains isolated from various environments. We performed a whole genome-based analysis of population structure, which revealed that *P. biforme, P. camemberti* and *P. fuscoglaucum* formed separate and specific genetic clusters, *P. biforme* and *P. camemberti* being sister clades (Figure 1A). *Penicillium camemberti* and *P. biforme* were both specific to the cheese environment, whereas *P. fuscoglaucum* was mostly found in natural environments. These findings suggest an ancient domestication event separating *P. biforme* from *P. fuscoglaucum* and a more recent domestication event separating the *P. camemberti* clonal lineage from *P. biforme.* Consistent with this scenario, we found evidence of phenotypic adaptation to cheese-making in *P. biforme* relative to *P. fuscoglaucum*, with a whiter color, faster growth on cheese medium under cave conditions and lower levels of toxin production. We also reveal the diversification of *P. camemberti* into two varieties, *P. camemberti* var. “*camemberti”* and var. *“caseifulvum”*, also displaying genetic and phenotypic differentiation, and maybe used for the production of different cheese types.

## Results

### *Penicillium camemberti, P. biforme* and *P. fuscoglaucum* each form separate specific genetic clusters

We collected and sequenced the genomes of 61 strains with Illumina technology, including 36 strains isolated from the rinds of various types of cheese (e.g., tommes, Camembert, blue cheeses), 11 from other food products (e.g. dry sausages, bread), and 14 from non-food environments (e.g., wood or leaf litter), which were attributed by their collectors to the species *P. camemberti, P. biforme, P. commune* or *P. fuscoglaucum* (Table S1). We resequenced the reference genome of *P. camemberti* (LCP06093, also known as FM013, initially sequenced with 454 technology [26]) with PacBio long-read technology and used the ca. 35Mb assembly obtained for mapping. We identified 392,072 SNPs across all strains (Supplemental Data 1). Principal component analysis (PCA; Figure 2A) and neighbor-net (splitstree) analysis (Figure 2C) identified three genetic clusters, corresponding to *P. camemberti* (*n*=19)*, P. biforme* (*n*=28) and *P. fuscoglaucum* (*n*=14). NGSadmix identified the same three genetic clusters at *K*=3 (Figure 2D), the *K* value at which the structure was the strongest and clearest. The genetic differentiation between *P. camemberti, P. biforme* and *P. fuscoglaucum* was further confirmed by the high values of fixation (*F_ST_*) and absolute divergence (*d*_*xy*_) indices (Table S2).

**Figure 2:**
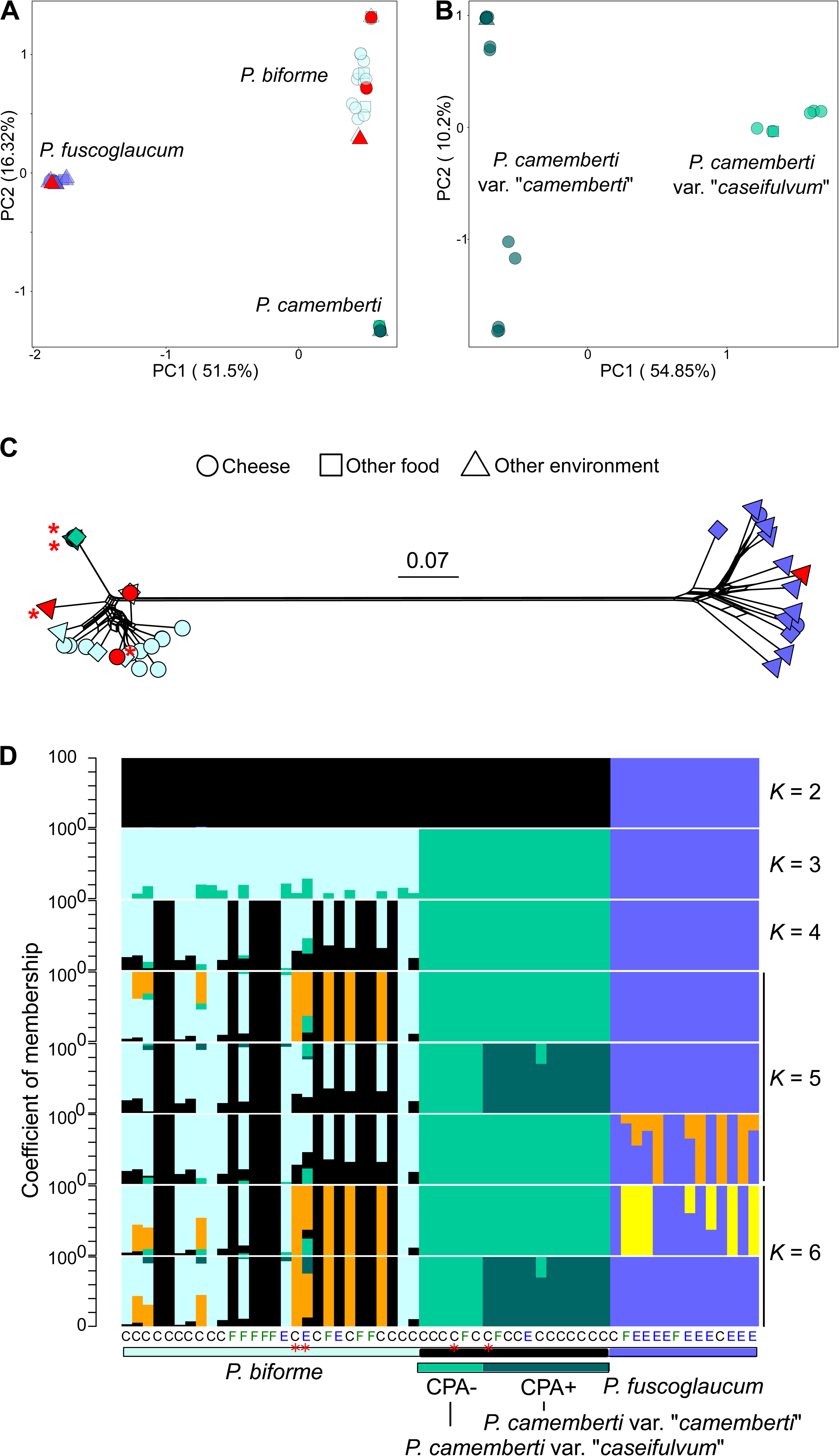
Population structure of 61 strains from the *Penicillium camemberti* species complex, based on whole-genome data. A) Principal component analysis (PCA) based on the 61 strains, B) Principal component analysis (PCA) based on only *P. camemberti* strains (n = 19). C) Neighbor-net analysis based on SNP data. Cross-linking indicates the likely occurrence of recombination. Branch lengths are shown and the scale bar represents 0.07 substitutions per site. D) Population subdivision inferred from *K* = 2 to *K* = 6. Each column represents a strain and colored bars represent their coefficients of membership for the various gene pools. For all panels, genetic clusters are represented by the same colors: dark purple for *P. fuscoglaucum,* light blue for *P. biforme,* dark green for *P. camemberti* var. “*camemberti*” and light green for *P. camemberti* var. “*caseifulvum*”. The strains identified as *P. commune* (non-monophyletic group) by collectors are indicated in red. For PCA and neighbor-net analysis, symbols correspond to the environment of collection: circles for cheese, triangles for dried sausages and squares for other environments. On the structure analysis, letters below the plots indicate the origin of the strains: C for cheese, F for food strains other than cheese and E for other environments. Red asterisks indicated the type strains of *P. biforme, P. camemberti and P. caseifulvum.* See also Table S1, S2, Figures S1, S2, Data S3 and S4.

A recent study described rapid phenotypic change from green-gray to white in a “wild *P. commune*” strain [39], interpreting these changes as evidence that domestication can occur within four weeks. However, we found that this *P. commune* strain actually belonged to the cheese *P. biforme* clade, being genetically almost identical to strains found in commercial cheeses (Figure S1).

The strains present in public collections under the name “*P. commune”* did not form a single cluster nor even a monophyletic clade (red shapes in Figures 2A, 2C and 3A); indeed, the *P. commune* strains isolated from natural environments clustered with *P. fuscoglaucum* strains whereas the *P. commune* strains isolated from cheese clustered with *P. biforme* strains, despite *P. fuscoglaucum* and *P. biforme* not being sister clades (Figure 3). The *P. commune* neotype strain (LCP05531) and the *P. biforme* type strain (LCP05529) belonged to the same clade (Figures 2C-D and Figure 3), further indicating that *P. commune* and *P. biforme* overlap. In contrast, the *P. caseifulvum* type strain (LCP05630) and the *P. camemberti* type strain belonged to different clusters at *K* values ≥ 5 in the population structure analysis (red stars on Figure 2C), suggesting that they do correspond to different genetic clusters.

**Figure 3:**
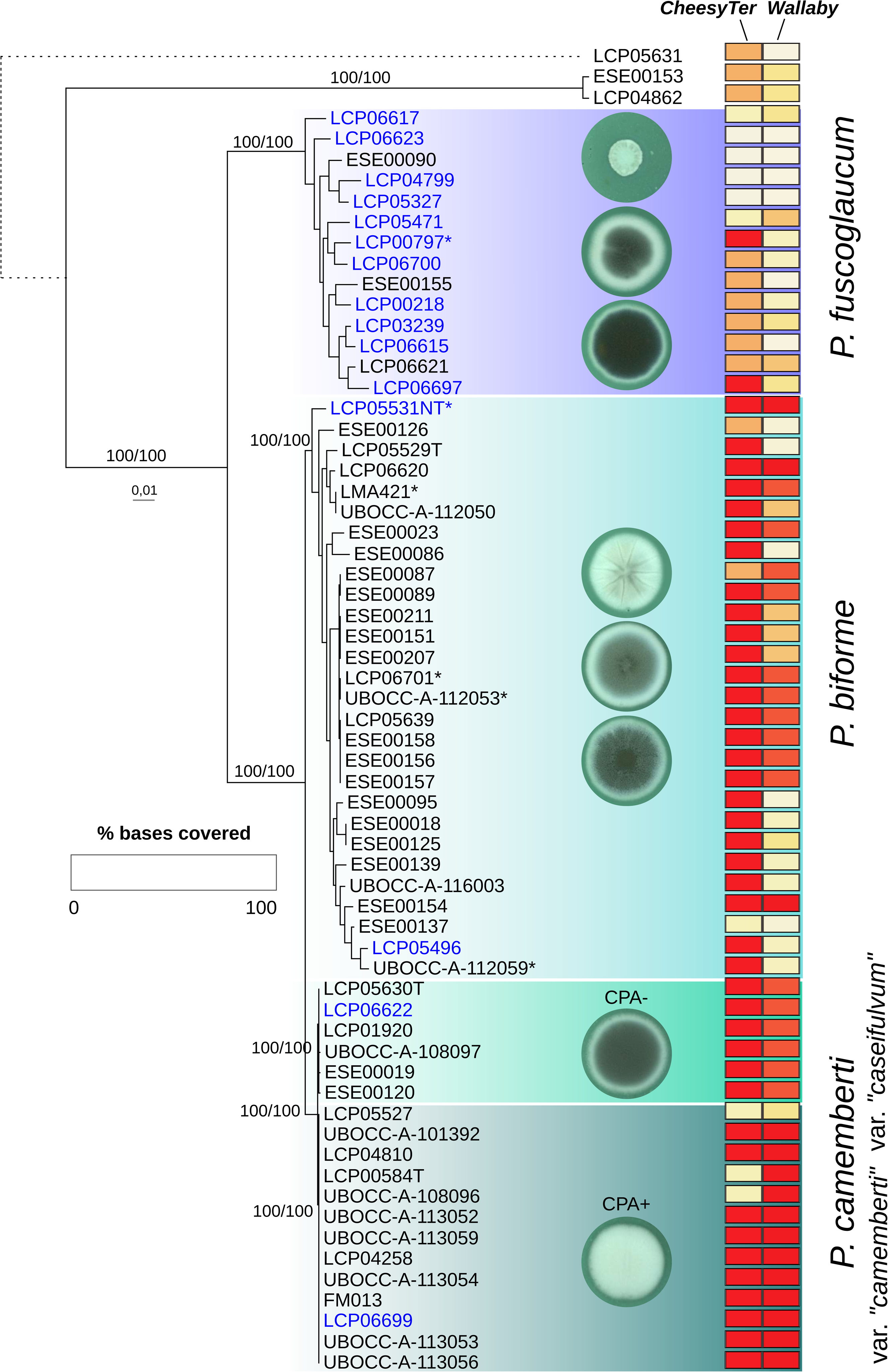
Phylogenetic relationships and presence/absence of the *Wallaby* and *CheesyTer* horizontally transferred regions in the 61 studied strains of the *Penicillium camemberti* species complex and the three outgroups. Maximum likelihood tree for the 64 strains based on 1,124,983 SNPs (586,131 parsimony informative sites). Branch supports, represented as SH-aLRT/ultrafast bootstrap in percentage, are only shown for clades but were always > 75%. The strain name color indicates the environment of collection, black for food and blue for other environments. Asterisks after strains indicate strains previously named *P. commune.* On the right panel, heatmap representing the percentage of bases covered in *Cheesyter* and *Wallaby* regions. See also Table S1, S2, Figures S1, S2, S3, Data S1, S3 and S4.

*Penicillium camemberti* was found only in food (cheese, sausage, food waste or animal feed; Figure 2). The non-cheese strains were not differentiated from cheese strains. Only three of the 28 strains in the *P. biforme* cluster were isolated from environments other than food: the atmosphere, ice and leaf litter. These three strains were not genetically differentiated from those isolated from food (bread, cheese or sausage) and may therefore be feral strains, i.e. escaped from food. The *P. fuscoglaucum* cluster included three strains isolated from food environments (two strains from cheese and one from sausage) and 13 strains from natural environments, and was thus likely to constitute a genuine wild population. The three food strains were not differentiated from the wild strains.

*Penicillium fuscoglaucum* displayed the highest genetic diversity level, followed by *P. biforme,* as indicated by the number of SNPs, π and Watterson’s *θ* (Table S2), as well as strain scattering in PCA and neighbor-net network (Figures 2A and 2C). The very low genetic diversity detected within *P. camemberti* suggests that it represents a clonal lineage (Table S2, Figures 2A and 2C). The average nucleotide identity between *P. camemberti* and *P. biforme* was very high (99.8%) and similar to the 99.5% identity between the domesticated mold *Aspergillus oryzae* and its wild relative *A. flavus* [40].

The long branches and the lack of reticulation observed in the neighbor-net analysis further confirmed the clonality of *P. camemberti*, contrasting with the footprints of recombination detected within both *P. biforme* and *P. fuscoglaucum* (Figure 2C). In *P. fuscoglaucum,* abundant reticulation was observed, right to the branch edges (Figure 2C), reinforcing the view that *P. fuscoglaucum* corresponded to a sexual population thriving in natural environments, while strains of *P. camemberti* and *P. biforme* used in the food industry are replicated clonally.

In heterothallic fungi, sexual reproduction occurs only between two haploid individuals carrying different mating-type alleles (MAT1-1 and MAT1-2 in heterothallic ascomycetes). A lack of sexual reproduction leads to relaxed selection on the mating-type genes and a departure from a balanced ratio between mating types. Neither mating-type allele presented any evidence of loss-of-function mutations or SNPs in any of the genomes. We found a significant departure from the 1:1 ratio in *P. biforme* (*χ*2 = 14.29; df = 1; P < 1e-3), with MAT1-2 strains predominating, but not in *P. fuscoglaucum* (*χ*2 = 1.14; df = 1; P = 0.29). All 19 *P. camemberti* strains carried the MAT1-2 allele, providing further evidence of the clonality of *P. camemberti* [41,42]. The observed clonality in *P. camemberti* may be explained by the recent selection of a white mutant and its use in industrial cheese production [43], whereas *P. biforme* is more widely used in the production of artisanal cheeses and farmers may make use of diverse local strains.

### Two genetic clusters within *Penicillium camemberti*

We found two genetic clusters within *P. camemberti*, as indicated by the population structure analysis implemented in NGSadmix at *K* = 5, the PCA and the neighbor-net without *P. fuscoglaucum*, as well as the PCA with only *P. camemberti* (Figures 2B, 2D and S2). The two *P. camemberti* genetic clusters displayed very few shared SNPs (13% of the 19,510 SNPs, Table S2), but a very high nucleotide identity (99.998%). The two genetic clusters within *P. camemberti* separated strains isolated from soft cheeses, such as Camembert, and strains isolated from other kinds, such as Rigotte de Condrieu and Saint Marcellin. The *P. caseifulvum* type strain (red star on Figure 2D) clustered with *P. camemberti* strains isolated from non-soft cheeses. The genetic cluster including the *P. caseifulvum* type strain will be referred to hereafter as *P. camemberti* var. “*caseifulvum*”, and the genetic cluster including the *P. camemberti* type strain as *P. camemberti* var. “*camemberti*”. The nucleotide diversity was very low within both varieties (Table S2, Figure 2B). We designed two pairs of primers to identify by a single PCR reaction the *P. camemberti* variety, each yielding a specific amplicon size.

The population structure analysis also suggested further genetic subdivision within each of *P. fuscoglaucum* and *P. biforme*; these subdivisions however did not correspond to the environment from which the isolate was obtained, type of cheese, or to a strong subdivision in the PCAs or the neighbor-nets (Figures 2 and S1).

### Presence/absence polymorphism of the two horizontally transferred regions likely to be advantageous in cheese

The PacBio-based genome assembly of the FM013 reference genome showed that *Wallaby* formed a single block (scaffold 18, positions 102,711 to 538,265). Two genes in *Wallaby* have been highlighted as putatively important for cheese-making. One, encoding the *Penicillium* antifungal protein (PAF) [26], was absent from FM013, whereas the other, *Hce2,* encoding a protein with antimicrobial activities, was present. All *P. camemberti* var. “*camemberti*” strains harbored the same *Wallaby* region as FM013 (Figures 3 and S3b), whereas strains from the *P. camemberti* var. “*caseifulvum*” lineage carried fragmented and duplicated segments of the *Wallaby* fragment present in FM013, with *Hce2* being systematically present. We found *Wallaby* fragmented with some duplicated regions in 18 *P. biforme* strains and completely absent from 10 strains (Figure 3 and S3b). *Wallaby* appeared to be even more fragmented in *P. fuscoglaucum* strains, always lacking *Hce2* but in one strain, and completely absent in nine strains. Fragments of *Wallaby* were found in some non-cheese *P. fuscoglaucum* strains. Even in *P. fuscoglaucum*, no genetic variability could be detected within the *Wallaby* fragments, indicating that they have also likely been acquired by horizontal gene transfer in this species. So far, we found no species in which *Wallaby* showed genetic variability, indicating that we have not identified the donor species.

We found no strict association between *Wallaby* and the 57 kb *CheesyTer* region, the horizontally transferred region thought to be beneficial for growth on cheese [27]. In *P. camemberti,* 16 of 19 strains carried the whole *CheesyTer* region, with an identity of 100% (Figures 3 and S3a, scaffold 17, positions 690,636 to 751,978 in FM013). The other three *P. camemberti* strains, belonging to the *P. camemberti* var. “*camemberti*” cluster, completely lacked *CheesyTer* and an 80 kb-downstream fragment. These three strains were isolated from Camembert cheeses around 1905, suggesting that the horizontal transfer of *CheesyTer* to *P. camemberti* occurred after 1905, in a single strain that has been since clonally cultured. In *P. biforme,* 25 of 28 strains carried *CheesyTer*, with an identity >99.8% between strains. The identity between *P. biforme* and *P. camemberti* was also >99.8%, except in the 5 kb region at the start of *CheesyTer,* in which the similarity between the two species dropped to 90%, mainly due to C:G to T:A mutations in *P. camemberti,* corresponding to the repeat-induced point (RIP) mutations specific to fungi occurring in repeated sequences during sexual reproduction [44]. As *P. camemberti* was found to be clonal and no RIP footprints were detected within the rest of *CheesyTer*, these findings suggest that *CheesyTer* was transferred to an ancestor of the two *P. camemberti* varieties with this RIPed sequence already present. The two *CheesyTer* genes involved in lactose metabolism [27], one encoding a lactose permease and the other a β-galactosidase, were present in all strains carrying at least one part of *CheesyTer* (Figure 3 and S3a).

### Looking for genomic footprints of adaptation in *P. camemberti* and *P. biforme*

Not many genomic selection scans could be performed given the high level of clonality in *P. camemberti*, as any selective sweep will hitchhike the whole genome. We investigated whether some genomic regions showed greater differentiation or higher diversity than the genomic background when comparing *P. biforme* to either *P. camemberti* or *P. fuscoglaucum*. No particularly striking genomic island of differentiation based on *d*_*xy*_ or diversity based on π and Watterson’s *θ* could be identified (Data S3, S4). The genes within windows in the 1% highest *d*_*xy*_ were not enriched in any particular function but contained significantly less genes with predicted functions than the rest of the reference genome, suggesting that these regions contained rapidly evolving genes.

### Phenotypes better suited for cheese production in *P. camemberti* and *P. biforme* than in the closely related wild *P. fuscoglaucum*

We performed a set of laboratory experiments to determine whether cheese strains had evolved traits beneficial for cheese production not present in the closely related wild *P. fuscoglaucum* lineage occurring in other environments. In each of *P. camemberti* and *P. biforme,* we considered as a single group the strains from cheese and other food types (e.g. dry sausage, bread) because they were not genetically differentiated and the same strains are sold by producers for cheese and dry sausage maturation.

We first investigated whether *P. camemberti* and *P. biforme* had acquired traits enabling them to grow more rapidly than *P. fuscoglaucum* in cheese-making conditions. Rapidly growing fungi can be beneficial for cheese-making, as they are more capable of excluding spoiler bacteria, yeasts and molds [10] and promote faster cheese maturation. Cheese maturation occurs in dark caves at a low temperature and high humidity. We therefore grew 61 strains on various media (unsalted cheese, salted cheese, minimal and malt media), in the dark and at different temperatures and humidity levels (25°C at ambient relative humidity, 10°C at 98% humidity or 8°C at 100% humidity), and we measured colony radial growth after 10 days. The non-parametric Kruskal-Wallis test and the ANOVA showed no significant effect on growth of the substrate from which the isolate was obtained (i.e. food versus non-food). We found a significant effect on growth of the culture media, species, temperature/humidity conditions and the *P. camemberti* variety. The ANOVA further showed significant effects on growth of the interactions between culture medium and species on the one hand and between culture medium and temperature/humidity on the other hand (Data S1A and S2A, Figure 4A). However, the significance of the interaction between culture medium and temperature/humidity conditions (Data S1A and S2A) reflected a very slow growth at low temperatures, making it difficult to detect differences between media. Although the two *P. camemberti* varieties were genetically very closely related, *P. camemberti* var. “*caseifulvum*” grew similarly to *P. biforme* (post-hoc Tukey test *P* = 1), with greater radial growth on unsalted cheese and salted cheese, and slower growth on malt and minimal media than *P. fuscoglaucum*. By contrast, *P. camemberti* var. “*camemberti*” displayed weaker radial growth than any other lineage on all media (Figure 4A), but was fluffier (Figure 1B). Furthermore, *P. camemberti* and *P. biforme* grew less rapidly than *P. fuscoglaucum* on minimal and malt media, indicating a disadvantage for growth in harsh conditions, as expected for cheese strains due to relaxed selection on functions useful only in wild environments. These findings demonstrated that *P. camemberti* var. “*caseifulvum*” and *P. biforme* have acquired rapid radial growth in cheese which was likely beneficial for fast maturation. *Penicillium camemberti* var. “*camemberti*” grew much less radially on cheese than any other lineage, but grew much more vertically (i.e. being fluffier), with the mycelium growing up to the lid of the Petri dishes (Figure 1B). *Penicillium camemberti* var. “*camemberti*” might have been selected for fast vertical growth to make soft cheeses with a fluffy aspect within a short maturation period. Short maturation is beneficial for production when using cow or goat milk available throughout the year, as is the case for Camembert cheeses. In contrast, ewe milk is only available a few months a year, so that Roquefort cheeses need to be stored for several months to be sold and consumed throughout the year. Such long storage can explain a selection for slow growth in the Roquefort *P. roqueforti* population before fridges were available [25], in contrast to a selection for fast growth in *P. camemberti* and *P. biforme*.

**Figure 4:**
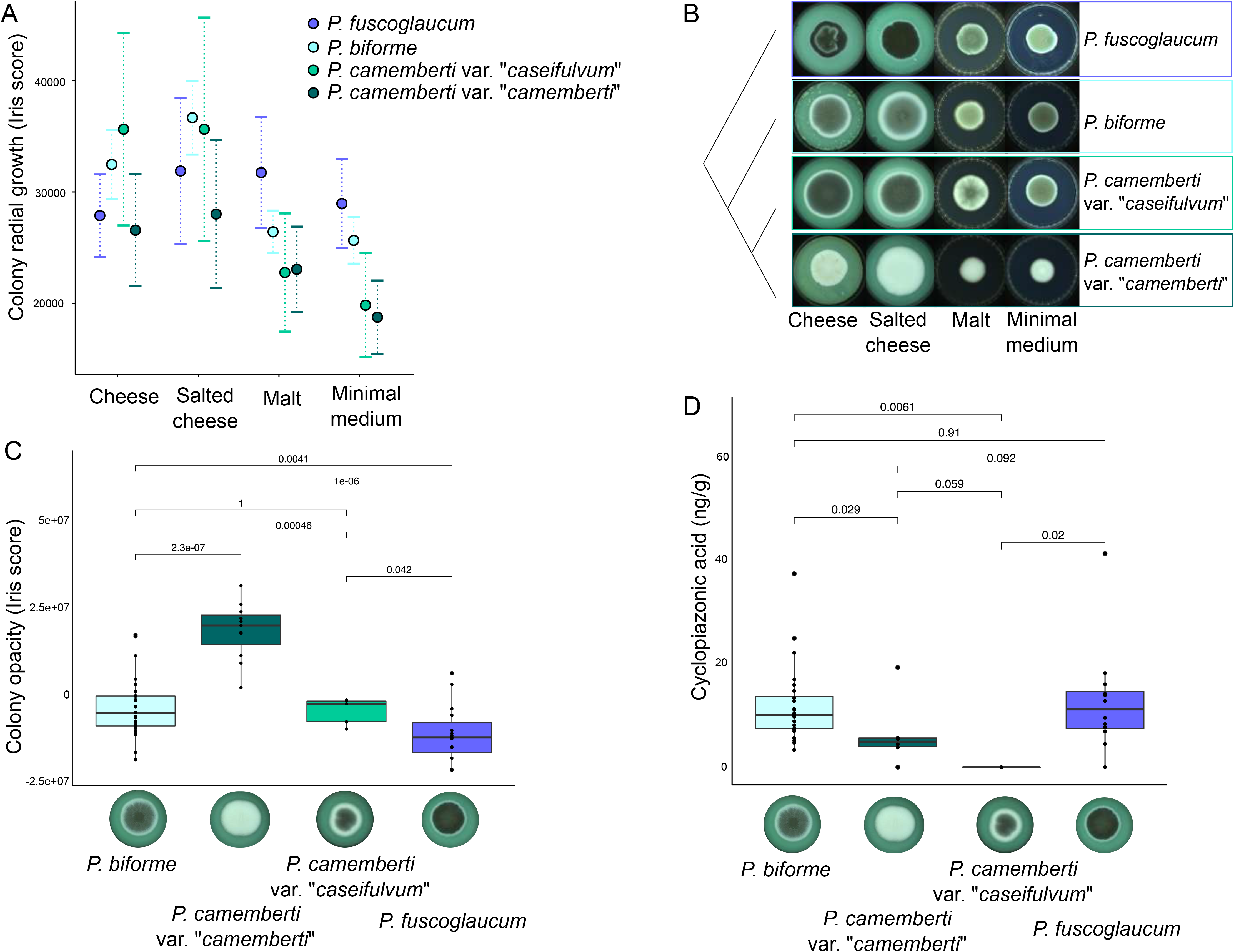
Phenotypic traits distinguishing the four lineages *Penicillium biforme, P. camemberti* var. “*caseifulvum*”*, P. camemberti* var. “*camemberti*” and *P. fuscoglaucum.* A) Mean radial growth of the four lineages on unsalted cheese, salted cheese, malt and minimal media. B) Pictures of colonies on unsalted cheese, salted cheese, malt and minimal media; *P. fuscoglaucum*: LCP05471; *P. camemberti* var. “*camemberti*”: UBOCC-A-113052; *P. biforme*: ESE00211; *P. camemberti* var. “*caseifulvum*”: LCP06622. C) Difference in opacity between the four lineages on salted cheese at 25°C. D) Difference in production of cyclopiazonic acid between the four lineages on yeast extract sucrose. C-D) Horizontal lines of the boxplots represent the upper quartile, the median and the lower quartile. Each dot represents a strain and p-values are given for the tests of differences between the lineages (Wilcoxon test). The color indicates assignments to the four lineages as in other figures. See also Table S1, Data S1, S2.

High salt concentrations in cheeses prevent the growth of contaminants and cheese fungi may have adapted to such conditions. We found an effect of salt on *P. biforme* growth, with strains growing more rapidly on salted than unsalted cheese medium (post-hoc Tukey test *P* = 0.006) and more rapidly on salted cheese medium than *P. camemberti* (post-hoc Tukey test *P* = 0.0001). Salt had no effect on the growth of *P. camemberti* (post-hoc test *P* = 1) or *P. fuscoglaucum* (post-hoc test *P* = 1). There may have been stronger selection for salt tolerance in *P. biforme,* which is used to inoculate blue and goat cheeses, both being more salty than soft cheeses, such as Brie or Camembert [45], for which *P. camemberti* var. “*camemberti*” is used.

We investigated whether cheese lineages had become whiter, which can be more attractive to some consumers than gray-green mold on cheese, by comparing the opacity of lineages on cheese medium, as opacity increases with the brightness and fluffiness of a colony [46]. We found significant effects of species and variety (Figure 4C and Data S1B and S2B), with *P. camemberti* being significantly more opaque (i.e. brighter and fluffier) than other lineages, and *P. camemberti* var. “*camemberti*” even more so compared to var. “*caseifulvum*” (post-hoc Tukey test *P* = 0.03). This is consistent with the white and fluffy aspect of the crust of Camembert and Brie, made with *P. camemberti* var. “*camemberti*”, whereas *P. camemberti* var. “*caseifulvum*” was found here in cheeses with a grayer and less fluffy crust, such as Saint Marcellin or Rigotte de Condrieu (Figures 1 and 4B-C). The *P. camemberti* var. “*caseifulvum*” and *P. biforme* lineages did not differ significantly from each other (post-hoc Tukey test *P* = 1) and both were brighter than the wild blue-green *P. fuscoglaucum* (Figure 4B).

We also investigated whether cheese lineages produced smaller amounts of cyclopiazonic acid (CPA), which is cytotoxic to humans [37,47], than strains isolated from other environments. *Penicillium camemberti* has been reported to produce CPA on yeast extract sucrose (YES) medium and at very low, non-toxic concentrations in cheese [38]. None of the *P. camemberti* var. “*caseifulvum*” strains tested here produced CPA on YES (Figure 4D), consistent with findings for the type strain [48]. By contrast, *P. camemberti* var. “*camemberti*”*, P. biforme* and *P. fuscoglaucum* produced CPA, the highest levels being obtained with *P. biforme* and *P. fuscoglaucum* (Figure 4D, Data S1C and S2C). The CPA biosynthesis cluster in the genome appeared to be functional, with its six genes present, but not the *cpaR* gene encoding a regulatory protein (Figure 5). The only exceptions were the six *P. camemberti* var. “*caseifulvum*” strains, in which a 2 bp deletion in the *cpaA* gene led to a frameshift. The *cpaA* gene encodes a polyketide synthase/non-ribosomal peptide synthase responsible for the first step of the CPA biosynthetic pathway so that a non-functional protein probably prevents CPA production on all substrates. Humans have often selected fungal strains unable to produce harmful toxins for use in food; *Aspergillus oryzae* strains used to ferment Asian food products, which do not produce aflatoxins, whereas its wild relative, *A. flavus,* does [49]. The *P. roqueforti* non-Roquefort population was found unable to produce mycophenolic acid due to a 174 bp deletion in the *mpaC* gene [50].

**Figure 5:**
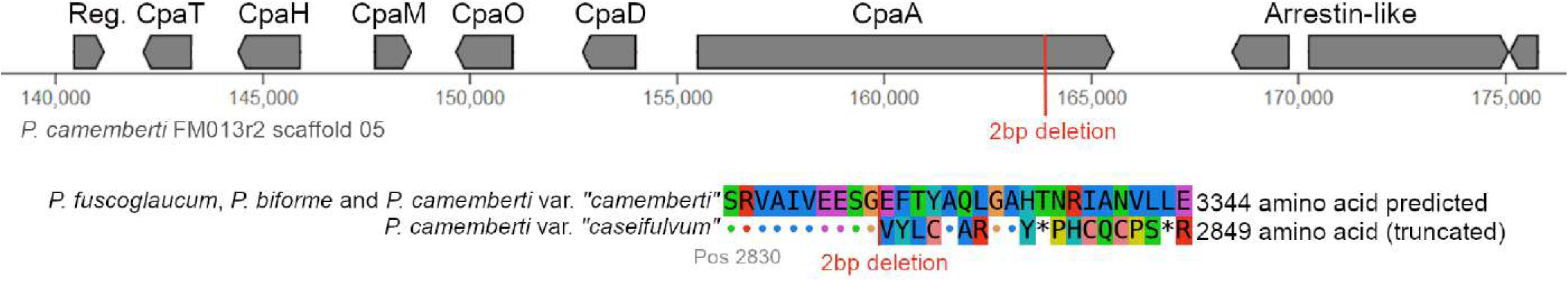
Cyclopiazonic acid (CPA) biosynthesis cluster in the *Penicillium camemberti* species complex. A) Schematic representation of the CPA biosynthesis cluster in the *P. camemberti* var. “*camemberti*” reference genome FM013. B) Amino-acid alignment of the *CpaA* gene showing a 2bp deletion in strains belonging to the *P. camemberti* var. “*caseifulvum*” genetic cluster, leading to a truncated protein. See also Table S1, Data S1, S2.

Cheese is a nutrient-rich environment in which many microorganisms can thrive, including undesirable food spoilage organisms. We therefore investigated whether the cheese lineages were better able to outcompete challengers. We allowed strains of *P. camemberti* var. “*camemberti*” (*n*=3), *P. camemberti* var. “*caseifulvum*” (*n*=2) or *P. biforme* (*n*=10) to grow as lawns on the surface of the cheese medium, which were inoculated 24 h later with a single spot, in the middle of the Petri dish, using a challenger: *Geotrichum candidum* (*n*=5)*, P. biforme* (*n*=6)*, P. fuscoglaucum* (*n*=5), or *P roqueforti* (*n*=12, three strains from each of the four known genetic clusters [25]). The species and the variety used as the lawn had significant effects on the growth of the challenger (ANOVA and non-parametric Kruskal-Wallis tests, Data S1D and S2D, Figure 6). The two *P. camemberti* lineages prevented the growth of all challengers more effectively than *P. biforme*, and *P. camemberti* var. “*camemberti*” was the most effective. We observed significant differences between the species used as challengers, with *G. candidum* having the highest growth differential on the various lawns, growing less well on a *P. camemberti* var. “*camemberti*” lawn than other lawns. *Geotrichum candidum* is a fungus that is also present on the surface of soft cheeses. The exclusion effect may be mediated by the fluffy morphology, resulting in the fungus occupying more space and using more resources, and/or by biochemical interactions.

**Figure 6:**
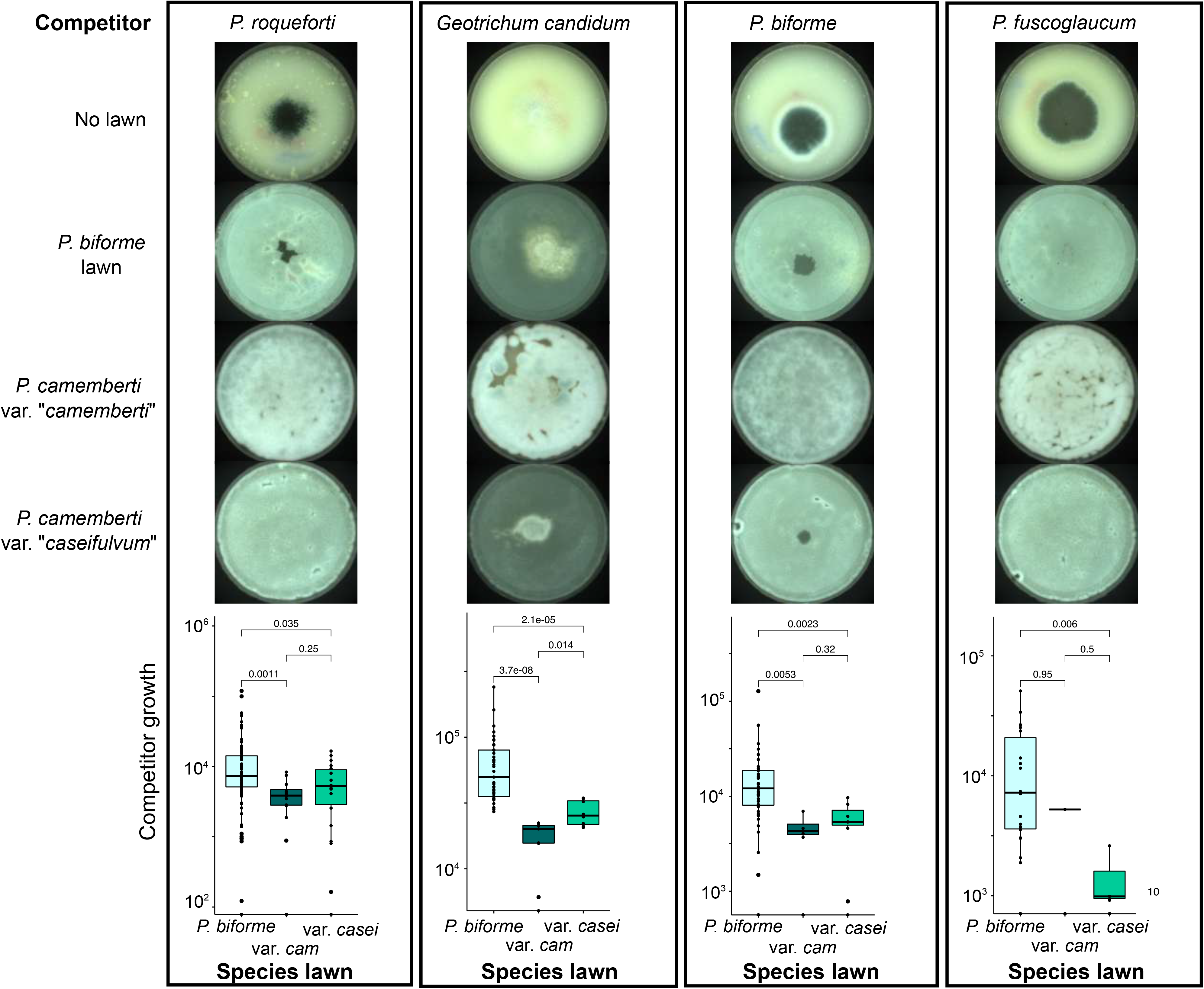
Competitive abilities of *Penicillium biforme, P. camemberti* var. “ *caseifulvum*” and *P. camemberti* var. “*camemberti*” against *Geotrichum candidum, P. biforme, P. fuscoglaucum* and *P. roqueforti* competitors. Top: Pictures of *P. roqueforti* FM164, *Geotrichum candidum* ESE00163, *P. biforme* ESE00154 and *P. fuscoglaucum* ESE00221 on no lawn (first line), *P. biforme* ESE00086 lawn (second line), *P. camemberti* var. “*camemberti*” LCP04810 (third line) and *P. camemberti* var. “*caseifulvum*” ESE00019 (fourth line) lawns, on salted cheese. Bottom: Boxplots representing the differences of growth abilities of the competitors on different species lawn. Horizontal lines of the boxplots represent the upper quartile, the median and the lower quartile. Each dot represents a strain and p-values are given for the tests of differences between the lineages (Wilcoxon test). The color indicates assignments to the three lineages as in other figures. Abbreviations: *P. camemberti* var. “*camemberti*”: var. *cam; P. camemberti* var. “*caseifulvum*”: var. *casei.* See also Table S1, Data S1, S2.

## Discussion

Whole-genome sequences revealed that *P. biforme, P. camemberti* and *P. fuscoglaucum* each formed a separate genetic cluster, with *P. biforme* and *P. camemberti* being sister clades. Strains identified as *P. commune* corresponded to either *P. biforme* or *P. fuscoglaucum*, being thus non-monophyletic (Figure 1A), so this name should not be used any longer. Furthermore, considering all *“P. commune”* strains to be wild, as it is currently common in the literature, leads to wrong inferences about trait evolution [39], as some *“P. commune”* strains belong to a domesticated genetic cluster. We found that *P. camemberti* and *P. biforme* were sister clades specific to the cheese environment, whereas the more distantly related clade *P. fuscoglaucum* was found mostly in natural environments. These relationships and the lower diversity of *P. biforme* than *P. fuscoglaucum*, with even lower levels of diversity in *P. camemberti*, suggest that an ancient domestication event led to the divergence between *P. biforme* and *P. fuscoglaucum* and that a much more recent domestication event led to the differentiation between *P. camemberti* and *P. biforme*. We used here the *P. biforme, P. camemberti* and *P. fuscoglaucum* Latin species names as they are commonly used and corresponded to differentiated and monophyletic clades based on genome-wide information. We did not address here the question of species status or definition, as they are notoriously difficult to elucidate among closely related fungal groups, especially in clonal ascomycetes [51,52].

We found evidence of phenotypic adaptation to cheese-making in *P. biforme* relative to the wild *P. fuscoglaucum* clade, with a whiter color, faster growth on cheese medium under cave conditions and lower levels of toxin production. These signs of domestication were even more marked in *P. camemberti* than in *P. biforme*. We observed a similar evolution of traits as in populations of the domesticated blue-cheese fungus *P. roqueforti*, which grows more rapidly on cheese medium, is more competitive against contaminant microorganisms, and grows less well under harsh conditions than wild strains [25,27]. Such convergent evolution under similar selective pressures suggests that evolution may repeat itself, as already reported in natural populations of *Anolis* lizards [53,54], three-spined sticklebacks [55] or Mexican cavefishes [56]. Phenotypic convergent evolution has been also reported in domesticated organisms, particularly in crop plants, for the loss of seed shattering, minimization of seed dormancy and increase in seed size and number [57,58].

We identified two genetically differentiated *P. camemberti* varieties, *P. camemberti* var. “*camemberti*” and *P. camemberti* var. “*caseifulvum*”, with contrasting phenotypic features, used in the production of different kinds of cheese. The *P. camemberti* var. “*camemberti*” strains were white and were isolated from Camembert or Brie. Their radial growth was slower but their mycelia were fluffier than the other *Penicillium* strains tested and they produced similar amounts of CPA compared to *P. biforme* strains. They excluded fungal competitors effectively, as previously suggested [59], probably due to their fluffy morphology, taking up the available space and monopolizing resources. *Penicillium camemberti* var. “*caseifulvum*” strains were gray-green, unable to produce CPA, and were isolated from cheeses other than Camembert, such as St Marcellin or Rigotte de Condrieu. They displayed more rapid radial growth than *P. camemberti* var. “*camemberti*” and grew similarly to *P. biforme* strains on cheese medium in cave conditions. The difference in terms of cheese type usage between the two varieties should be confirmed using more strains, but it seems to correspond well to the contrasted identified phenotypes, and in particular color and fluffiness. The existence of two genetically and phenotypically differentiated lineages with possibly different uses in cheese-making suggested that these lineages emerged as a result of different human selection pressures, as reported for the domesticated blue-cheese fungus *P. roqueforti* [25], wine-making yeasts [11,14], maize, rice, tomatoes, dogs, chickens and horses [1–6].

*Penicillium camemberti* var. “*camemberti*” is the emblematic fungus used to inoculate soft cheeses, such as Camembert and Brie. According to the technical specifications for Brie de Meaux and Brie de Melun PDOs and Camembert, the crust of the cheese must be white and fluffy, and inoculation with *P. candidum* (a synonym of *P. camemberti*) is even specified for Brie de Meaux. These specifications are recent (20th century) and seem to have had a negative impact on the diversity of the fungi used for making these kinds of cheeses, as a single clonal lineage now predominates. The genetic diversity of *P. roqueforti* has also been greatly reduced by recent industrialization [25,28], although PDO specifications that local strains must be used have protected diversity to some extent in Roquefort cheeses. Camembert and Brie cheeses were gray-green until the middle of the 20th century [31,33,43]. First mentions of Brie cheeses date from middle ages [31]. These historical records are consistent with our findings, which suggest a first domestication event leading to the emergence of the gray-green mold *P. biforme,* subsequently followed, around 1900, by the domestication of *P. camemberti,* with the selection of two different varieties displaying more marked signs of domestication than *P. biforme*.

These findings have industrial implications, as they reveal the existence of three closely related but different lineages that have evolved traits beneficial for cheese-making, with different phenotypic traits selected according to usage. This study should foster further research, as it would be interesting to assess other important traits, such as volatile compound production, and the efficiencies of lipolysis and proteolysis. Our findings raise questions about the use of limited number of clonal strains for cheese-making, which tends to lead to degeneration, limiting the possibilities for further improvement, which is currently a major concern in the agrofood sector [60], despite the great geneticist Nikolai Vavilov known for having identified the centers of origin of cultivated plants long ago highlighting the importance of genetic diversity in domesticated organisms for variety improvement and diversification [61].

## Acknowledgments

We thank everyone who sent us cheese crusts, Steve Labrie for the LMA strain, and Joëlle Dupont for strains from the National Museum of Natural History in Paris.

This work was supported by the Fungadapt ANR-19-CE20-0002-02 ANR grant to TG, JR, EC, MC and a Fondation Louis D. grant from the Institut de France to TG.

## Author contributions

J.R. and T.G. designed and supervised the study, and obtained funding. J.R., E.D., A.S. and S.LP. generated the data. J.R., R.C.RdlV. and B.B. analyzed the genomes. J.R., E.D. and A.S. performed the experiments. M.C., E.C. and E.P. performed the mycotoxin experiment and contributed to CPA biosynthetic gene cluster analyses. J.R. and E.D. analyzed the data from the laboratory experiments. J.R., and T.G. wrote the manuscript, with contributions from all the authors. J.R., T.G. and R.C.RdlV. prepared the final version of the manuscript.

## STAR Methods

### Resource availability

#### Lead contact

Further information and requests for resources and strains with ESE numbers should be directed to and will be fulfilled by the Lead Contact, Jeanne Ropars.

#### Materials Availability

Strains used in this study are publicly available: strains with LCP identifiers are from the public collection of the National Museum of Natural History in Paris; strains with UBOCC identifiers are from the public collection of the Université de Bretagne Occidentale Culture Collection; other strains with ESE numbers are conserved at the Ecology, Systematics and Evolution laboratory in Orsay.

#### Data and Code Availability

Raw reads generated in this study have been deposited to NCBI under the project number PRJNA655754.

The dataset corresponding to phenotype measurements is available in Data S1.

### Experimental model and subject details

We analyzed strains isolated from 37 cheeses from five countries around the world (e.g. Camembert, Saint Marcellin, tomme, Sein de Nounou). We also collected strains from dried sausages (*n* = 8) and moldy food (e.g. bread, *n* = 4). Twelve were isolated from other substrates than food. Spores were sampled from the food sources and spread on Petri dishes containing malt-agar medium, which were then incubated for five days at 25°C. A total of 41 strains were obtained from public strain collections (the Laboratoire de Cryptogamie (LCP strains) at the National Museum of Natural History in Paris (France), the Laboratoire Universitaire de Biodiversité et Ecologie Microbienne (LUBEM, UBOCC strains) in Plouzané (France) and the Westerdijk Fungal Biodiversity Institute (CBS strains) in Utrecht (The Netherlands)). The LMA421 strain was kindly given by Steve Labrie (University of Laval, Canada). We obtained strains from natural environments (e.g. wood or natural cave walls) from eight different countries. Detailed information about the strains used in this study can be found in the Table S1. For each strain, single-spore cultures were generated by a dilution method, to ensure that only a single haploid genotype was cultured for each strain. We checked the species identification of all strains by Sanger sequencing of the ß-tubulin gene (primers Bt2a - GGTAACCAAATCGGTGCTGCTTTC and bt2b – AACCTCAGTGTAGTGACCCTTGGC [62]) and the PC4 microsatellite flanking regions [35] (primers PC4F – CAAGCTGGCCGATAACCTG and PC4R – CCATCCGCTTGATTTCTCCT), to distinguish between *P. biforme* and *P. camemberti*. All the strains were added to the ESE (Ecology Systematics Evolution Laboratory) collection, under an ESE accession number, and are available from the corresponding collections (ESE, LUBEM, MNHN).

### Method details

#### DNA extraction

For each strain, we used the Nucleospin Soil Kit (Macherey-Nagel, Düren, Germany) to extract DNA from fresh haploid mycelium grown for five days on malt agar.

#### Genome sequencing, assembly and mapping

Sequencing was performed with Illumina HiSeq 2500 paired-end technology (Illumina Inc.), with a mean insert size of 400 bp, at the INRA GenoToul platform, to obtain 10-50X coverage (Table S1). In addition, the genome of the *P. camemberti* LCP06093 (also known as FM013) reference strain, initially sequenced with 454 technology [26], was resequenced at 34x using the PacBio’s long-read technology, assembled with Canu 1.8 [63] and polished with pilon 1.23 [64]. The PacBio assembly and reads for all samples have been deposited at the NCBI Sequence Read Archive under BioProject ID PRJNA655754.

Reads were trimmed with Trimmomatic [65] (cut adapter, options PE, LEADING:3 TRAILING:3 SLIDINGWINDOW:4:25 MINLEN:36) and mapped onto the high-quality PacBio reference genome of LCP06093 with Bowtie2 [66] (options very-sensitive-local, --phred33 PL:ILLUMINA-X 1000). Mapped reads were filtered for PCR duplicates with picard tools (http://broadinstitute.github.io/picard) and realigned on the PacBio reference genome with GATK [67]. Single-nucleotide polymorphisms (SNPs) were called with GATK HaplotypeCaller, which provides one gVCF per strain (option -ERC GVCF). GVCFs were combined with GATK CombineGVCFs, genotyped with GATK genotype and hard filtered in accordance with GATK best practice workflow recommendations (https://software.broadinstitute.org/gatk/best-practices/).

#### Analysis of the mating-type genes

We looked for the mating type genes in *P. camemberti, P. biforme* and *P. fuscoglaucum* by extracting the region including the flanking genes *APN2* and *SLA2* from all BAM files.

#### Identification by a single PCR reaction the *P. camemberti* variety

We designed two pairs of primers to identify by a single PCR reaction the *P. camemberti* variety, each yielding a specific amplicon size. The forward primer is common to both reactions (Pcamcasei_F – CAAATCGAGGAATCACAAGTCT); Reverse primers sequences are Pcam_R – GAAATTCGATCTTGGGCAAA and Pcasei_R – GAGTAAGCATTGGCCTCGAC. The expected product size for *P. camemberti* var. “*camemberti*” isolates is of 776bp, whereas the expected product size for *P. camemberti* var. “*caseifulvum*” isolates is of 233bp.

PCR was performed in 30 μL reactions, using 15μL of template DNA (5ng/μL), 0.15 μL of Taq DNA polymerase (5 U/μL, mpbio medical), 3 μL of buffer with MgSO4 (25mM), 1.2 μL of dNTPs (10mM), 0.6 μL of each reverse primer, 1.2 μL of the forward primer and 15 μL of water.

Amplifications were performed on a Thermal cycler using a touchdown PCR, with 8 cycles of 30 s at 94°C, an initial annealing temperature of 60°C for 30 s gradually reduced by 2°C every cycle and 1min30 at 72°C. The program is followed by 24 cycles of 30 s at 94°C, 30 s at 53°C and 1min30 at 72°C. The program was ended by a 7 min extension step at 72°C.

PCR products were purified and sequenced by GENEWIZ, in one direction (forward).

Expected DNA sequence for *P. camemberti* var. “*camemberti*” is CCCCGGGAGGTTCCCTAGGCTGGCCGGGATGCTAGCTATATTAAAGAGGCCCCCGGCGATGCTAAAGCCAATTATCTTTAGTATATCGCATCGCTACCTTAGCATTTAGTTACTAATATTTGTTATATAACGGAATTTAACTCCTCGATTCTTTCTACCAAAATGATTGGCTAGTTAAGCTTAACGACAAGGTCGCCGACTTTAGCGCTAAGCAAGTCATCACCTTCGATATTGCTCTCACTTATTACAACCGCCCCTAGACTAGCCAATTACACACTAGGCTAAAAAGACCACTATATCTAAGTATTACGCTCAGATAGTCTAGGCTCCGTTATATATAGTGCGTGTTATAACATTACTACCATCATATTCGACAATATCATTATCCTCGCTCCCCGTATCGCCCTCGCTATCCGAGTCCATAAACCCAAAGAGAGCAAAGTATCAACACTATCAAGATCCACTACAATTCCACCATGATAGCGAAAAACCAACCATTAGCAATAAAAGGTCTATAGATCAGGGAGCTTCTCCAAGGTATCGCAAGGTACTGGATATACAACATAGACGACAAAATCACTAAGTCTTAACCACCCCTACGCTCCGACCCTAAATATATA GCGACTTCACCATTATGTGTAAAGGAGAAGAATATACTGTATATAGGTATATAGTTTGCCCAA and for *P. camemberti* var. “*caseifulvum*” CTGTTCTCCCCATATATGAAGAGAACAGTTTTGTCCTCCACTCCCACTATTTTTCCTGCGTCTCTATTTTATAATTTGTGATTACCTGATTCCCATCATGGCAGAAAAGATCTCCGGCAATATAGGTGTCGAGGCA.

#### Statistics of population genetics and genomic scans

Nucleotide diversity indices (π and Watterson’s *θ*) and genetic differentiation indices (*F*_*ST*_ and *d*_*xy*_) were calculated using the R package PopGenome [68] for the four genetic clusters *P. biforme, P. fuscoglaucum, P. camemberti.* var. “*camemberti*” and *P. camemberti* var. “*caseifulvum*”. π and Watterson’s *θ* were calculated per site. We also performed genomic scans along the 20 largest scaffolds of the *P. camemberti* PacBio reference using sliding windows of 75 kb and step sizes of 7.5 kb. Genome scans for π and Watterson’s *θ* were performed within each of the four genetic clusters (*P. biforme, P. fuscoglaucum, P. camemberti.* var. “*camemberti*” and *P. camemberti* var. “*caseifulvum*”). Genome scans based on *F*_*ST*_ and *d*_*xy*_ indices were performed between all possible pairs of the four clusters (*P. biforme, P. fuscoglaucum, P. camemberti.* var. “*camemberti*” and *P. camemberti* var. “*caseifulvum*”). We tested whether the genes belonging to windows with the 1% highest *d*_*xy*_ values were enriched in particular predicted functions (interpro). We performed gene enrichment analyses using Fisher exact tests (‘fisher.test’ function in the R package stats) corrected for multiple testing (‘p.adjust’ function in the R package stats with the false discovery rate, ‘fdr’, method). We also tested whether the number of genes with predicted functions within windows with the 1% highest *d*_*xy*_ values deviated from expectations based on the proportion in the full genome of the PacBio reference using a χ² test (Data S3, Data S4).

We calculated the average pairwise identity between species pairs, as well as the number of fixed, private and shared SNPs. When comparing two populations, private SNPs are sites that are polymorphic in only one population (e.g. A/C for individuals in population 1 and T for individuals in population 2), shared SNPs are polymorphic positions within the two populations (e.g. A/C in population 1 and A/T in population 2) and fixed SNPs are positions that are fixed in each population but different between populations (e.g. A for all individuals of population 1 and G for all individuals of population 2).

#### Genetic structure

We used the dataset of 392,072 SNPs to infer population structure. We inferred individual ancestry from genotype likelihoods based on realigned reads, by assuming a known number of admixing populations, ranging from *K* = 2 to *K* = 6, using NGSadmix from the ANGSD package [69]. A hundred independent runs were carried out for each number of clusters (*K)*. We identified clustering solutions in replicated runs for each *K* value and displayed population structure graphically using the R function ‘barplot’. We used the R package phangorn [70] for neighbor-net analyses, and the ‘prcomp’ function of R for principal component analysis (PCA).

#### Phylogenetic analyses

We inferred phylogenetic relationships among the 61 isolates using the dataset of 1,124,983 SNPs in a maximum likelihood framework using IQ-Ttree2 [71]. We also used *P. cavernicola* (LCP05631 strain) and *P. palitans* (ESE00153 and LCP04862 strains). The tree has been midpoint rooted. The best-fit model chosen using ModelFinder [72] according to BIC was TVM+F+ASC+R3. Branch supports are SH-aLRT [73] and ultrafast bootstrap support [74].

#### Laboratory experiments

##### Sampling and calibration

All experiments were performed on the whole collection of *P. camemberti* (*n* = 19)*, P. biforme* (*n* = 28) and *P. fuscoglaucum* (*n* = 14) strains, including 35 strains isolated from cheeses, 10 from other food environments (e.g. bread, sausage) and 14 strains isolated from environments other than food. Experiments were initiated with spore suspensions calibrated to 10^7^ spores/mL with a hemocytometer, under the constraint of the low rate of sporulation in *P. camemberti.*

##### Growth in different conditions and on different media

We investigated whether strains isolated from cheese displayed faster or slower radial growth than strains isolated from other environments when cultured on cheese or other substrates, and under conditions similar to those in maturation caves or other conditions. We prepared four different culture media: a cheese medium without salt, a salted cheese medium (17g/L, corresponding to a typical cheese), a malt medium and a minimal medium. The cheese media were produced from an unsalted drained cow’s milk cheese from Coubertin Farm in Saint Rémy-les-Chevreuse (France), as previously described [59]; we added five drops of blue food coloring to these media, to make it easier to distinguish white fungal colonies from the medium. The cheese media were rich in lipids and proteins, whereas the malt medium (20 g/L) was rich in carbohydrates. The minimal medium [75] contained only the trace elements necessary for fungal survival (i.e. iron sulfate, zinc sulfate, boric acid, magnesium chloride, copper sulfate, ammonium heptamolybdate, cobalt chloride, EDTA, sodium nitrate, potassium chloride, potassium phosphate and magnesium sulfate). All media were sterilized in an autoclave (121°C for 20 min for the malt and minimal media, 110°C for 15 min for the cheese media). Each 90 mm-diameter Petri dish was filled with 20 mL of the appropriate medium.

We allowed the fungi to grow in the dark, under three different conditions for each strain and each medium: 10°C with 98% humidity (Camembert cave conditions), 8°C with 85% humidity (cave conditions for other cheeses) and 25°C with ambient relative humidity (ambient conditions). We had 244 Petri dishes in total for each set of conditions, and we left the colonies to grow for 10 days. Images of the Petri dishes were obtained with a Scan 1200 from Interscience and analyzed with IRIS [46] for growth and color-opacity measurements. The colony size was estimated as the colony area in pixels. The opacity score was defined as the sum of the brightness values for all the pixels within the colony bounds; this score thus captures both the color and the three-dimensional colony growth (as opacity increases with fluffiness) and can be used as a proxy to assess the fluffiness of the fungal colony (Data S1, Data S2).

##### Mycotoxin production

We selected at least six strains from each genetic cluster for this experiment (Data S1). For measurements of mycotoxin production, we used 1 μL of calibrated spore suspension (10^7^ spores/mL) for each of the 61 strains to inoculate YES (yeast extract sucrose) agar medium buffered at pH 4.5 with phosphate-citrate buffer and characterized by a high C/N ratio to favor mycotoxin production as already described in [50]. Each culture was performed in triplicate for mycotoxin analyses as well as for fungal dry weight measurements. The plates were incubated at 25°C in the dark for 10 days and were then stored at −20°C until mycotoxin analysis.

For mycotoxin extractions [50], we homogenized the thawed samples with an Ultraturrax T25 (IKA, Heidelberg, Germany) before recuperating 4g aliquots. Then, 25 mL of acetonitrile (ACN) supplemented with 0.1% formic acid (v/v) was added, samples were vortexed for 30 sec followed by 15 min sonication. Extracts were then centrifuged for 10 min at 5000g and the recuperated supernatants were directly filtered through 0.2 μm PTFE membrane filters (GE Healthcare Life Sciences, UK) into amber vials. All samples were stored at −20°C until analyses.

Mycotoxin detection and quantification were performed using an Agilent 6530 Accurate-Mass Quadropole Time-of-Flight (Q-TOF) LC/MS system equipped with a Binary pump 1260 and degasser, well plate autosampler set to 10°C and a thermostated column compartment. Filtered samples (2μl) were injected into a ZORBAX Extend C-18 column (2.1×50mm and 1.8 μm, 600 bar) maintained at 35°C with a flow rate of 0.3 ml/min using mobile phase A (milli-Q water + 0.1% formic acid (v/v) and 0.1% ammonium formate (v/v) and mobile phase B (ACN + 0.1% formic acid). Mobile phase B was maintained at 10% for 4 min followed by a gradient from 10 to 100% for 16 min. Then, mobile phase B was maintained at 100% for 2 min before a 5 min post-time. Cyclopiazonic acid (CPA) was ionized in electrospray ionization mode ESI+ in the mass spectrometer with the following parameters: capillary voltage 4 kV, source temperature 325°C, nebulizer pressure 50 psig, drying gas 12 l/min, ion range 100-1000 m/z. CPA had a retention time of 15.3 min and was quantified using the [M+H]+ 337.1545 m/z ion and [M+Na]+ 359.1366 qualifier ion (ESI+). Other extrolites included in our internal database that were targeted (in ESI+ and ESI-modes) were andrastin A, citreoviridin, citrinin, ermefortins A & B, (iso)-fumigaclavin A, griseofulvin, meleagrin, mycophenolic acid, ochratoxin A, patulin, penitrem A, PR toxin, roquefortin C, sterigmatocystin.

We used a matrix matched calibration curve (R^2^ >0.99) for reliable mycotoxin quantification with final concentrations ranging from 10 to 10000 ng/ml. Method performance and mycotoxin determination was carried out as previously described in [50]. Specific mycotoxin production was expressed as ng per g fungal dry weight (Data S1, Data S2).

##### Competition

We inoculated salted cheese with 150 μL of a calibrated spore solution (10^7^ spores/mL), which we allowed to grow as a lawn. We used two *P. camemberti* var. “*caseifulvum*” strains (LCP05630 and ESE00019), three *P. camemberti* var. “*camemberti*” strains (FM013, LCP04810 and LCP05527) and 10 *P. biforme* strains (ESE00018, ESE00086, ESE00089, ESE00126, ESE00139, LCP05531, LCP06701, ESE00154, LCP06620 and LCP05496). After 24 h of growth, we deposited a single 20 μL drop of calibrated spore solution from a challenger species (10^7^ spores/mL) in the middle of the Petri dish. The challenger species used were *G. candidum* (*n*=4, ESE00163 and ESE00165, CBS11628 and VTTC4559)*, P. biforme* (*n*=6, ESE00126, ESE00089, ESE00018, ESE00154, LCP005531, LCP06701)*, P. fuscoglaucum* (*n*=5, ESE00090, LCP06621, LCP000797, LCP04799, LCP03239), and *P roqueforti* (*n*=12, three from each of the four genetic clusters described in [25]: LCP06036, LCP06037, LCP06128, LCP06043, LCP06098, LCP06064, FM164, LCP06137, LCP06131, LCP06173). The colony size of the challenger was calculated as the colony area in pixels (Data S1, Data S2).

### Quantification and statistical analysis

We performed analyses of variance (ANOVAs) followed by post-ANOVA Tukey’s HSD (honest significant difference) tests, with R. The data normality was improved with the bestNormalize R package [76]. For improving distribution normality, we used an ordered quantile (ORQ) transformation (a one-to-one transformation for vectors with an arbitrary distribution that generates normally distributed vectors) for diameter measurements and competition data and a Yeo-Johnson transformation for CPA (mycotoxin) data. Some residuals were still not normally distributed but ANOVA is known to be robust to deviations from normality [77]. When residues deviated from normality (diameter measurements for the growth and competition), we nevertheless also ran non parametric tests, i.e. Kruskal-Wallis tests, using R.

For ANOVAs, we used standard linear models in which all explanatory variables were discrete. The variables common for all analyses (with explained variables being growth, opacity, CPA and competition, respectively) were ‘species’, ‘*P. camemberti* variety’ and ‘substrate of origin’ (food versus non-food). The variables ‘medium’ and ‘temperature/hygrometry’ were explanatory variables specific to the growth analysis. The ‘challenger species’ variable was specific to the competition analysis. All variables and all interactions between them were implemented in the ANOVA, and non-significant interactions were subsequently removed before performing post-ANOVA Tukey’s HSD tests.

## Supplemental Table and Data Legends

**Data S1: Data for measured traits in *Penicillium camemberti* var. “*camemberti*”, *P. camemberti* var. “*caseifulvum*”, *P. biforme* and *P. fuscoglaucum*. Related to Figures 4, 5 and STAR Methods.**

A tab: growth; B tab: opacity; C tab: Cyclopiazonic acid production; D tab: competition.

**Data S2: Results of statistical analyses performed for testing differences in traits between *Penicillium camemberti* var. “*camemberti*”*, P. camemberti* var. “*caseifulvum*”*, P. biforme* and *P. fuscoglaucum*. Related to Figures 4, 5 and STAR Methods.**

Significant P-values are indicated with an asterisk and highlighted in gray. A tab: growth; B tab: opacity; C tab: Cyclopiazonic acid production; D tab: competition.

**Data S3: Genomic scans along the first 20 scaffolds of *d*_*xy*_, *F*_*ST*_, π and Watterson’s *θ*. Related to Figures 2 and 3A, Data S4 and STAR Methods.**

Dots show windows in the 1% highest *d*_*xy*_ between *Penicillium biforme* and *P. caememberti* var. “*caseifulvum*”, var. “*camemberti*” and *P. fuscoglaucum.*

**Data S4: Gene annotation within windows in the 1% highest *d*_*xy*_ between *Penicillium biforme* and *P. camemberti* var. “*caseifulvum*” (A tab), var. “*camemberti*” (B tab) and *P. fuscoglaucum* (C tab), between *P. fuscoglaucum* and *P. camemberti* var. “*caseifulvum*” (D tab) and var. “*camemberti*” (E tab). Related to Figures 2 and 3A, Data S3 and STAR Methods.**

## References

1. Warmuth, V., Eriksson, A., Bower, M.A., Cañon, J., Cothran, G., Distl, O., Glowatzki-Mullis, M.-L., Hunt, H., Luís, C., do Mar Oom, M., et al. (2011). European Domestic Horses Originated in Two Holocene Refugia. PLoS ONE 6, e18194.

2. Axelsson, E., Ratnakumar, A., Arendt, M.-L., Maqbool, K., Webster, M.T., Perloski, M., Liberg, O., Arnemo, J.M., Hedhammar, Å., and Lindblad-Toh, K. (2013). The genomic signature of dog domestication reveals adaptation to a starch-rich diet. Nature 495, 360–364.

3. Frantz, L.A.F., Schraiber, J.G., Madsen, O., Megens, H.-J., Cagan, A., Bosse, M., Paudel, Y., Crooijmans, R.P.M.A., Larson, G., and Groenen, M.A.M. (2015). Evidence of long-term gene flow and selection during domestication from analyses of Eurasian wild and domestic pig genomes. Nat. Genet. 47, 1141–1148.

4. Hufford, M.B., Xu, X., van Heerwaarden, J., Pyhäjärvi, T., Chia, J.-M., Cartwright, R.A., Elshire, R.J., Glaubitz, J.C., Guill, K.E., Kaeppler, S.M., et al. (2012). Comparative population genomics of maize domestication and improvement. Nat Genet 44, 808–811.

5. Decroocq, S., Cornille, A., Tricon, D., Babayeva, S., Chague, A., Eyquard, J.-P., Karychev, R., Dolgikh, S., Kostritsyna, T., Liu, S., et al. (2016). New insights into the history of domesticated and wild apricots and its contribution to Plum pox virus resistance. Mol. Ecol. 25, 4712–4729.

6. Doebley, J. (2004). The Genetics of Maize Evolution. Annual Review of Genetics 38, 37–59.

7. Wydooghe, E., Berghmans, E., Rijsselaere, T., and Soom, A.V. (2013). International breeder inquiry into the reproduction of the English bulldog. Vlaams Diergeneeskundig Tijdschrift 82, 38–43.

8. Perrier, X., Langhe, E.D., Donohue, M., Lentfer, C., Vrydaghs, L., Bakry, F., Carreel, F., Hippolyte, I., Horry, J.-P., Jenny, C., et al. (2011). Multidisciplinary perspectives on banana (*Musa* spp.) domestication. PNAS 108, 11311–11318.

9. Gladieux, P., Ropars, J., Badouin, H., Branca, A., Aguileta, G., de Vienne, D.M., Rodríguez de la Vega, R.C., Branco, S., and Giraud, T. (2014). Fungal evolutionary genomics provides insight into the mechanisms of adaptive divergence in eukaryotes. Molecular Ecology 23, 753–773.

10. Dupont, J., Dequin, S., Giraud, T., Le Tacon, F., Marsit, S., Ropars, J., Richard, F., and Selosse, M.-A. (2017). Fungi as a Source of Food. In Microbiology Spectrum.

11. Almeida, P., Gonçalves, C., Teixeira, S., Libkind, D., Bontrager, M., Masneuf-Pomarède, I., Albertin, W., Durrens, P., Sherman, D.J., Marullo, P., et al. (2014). A Gondwanan imprint on global diversity and domestication of wine and cider yeast *Saccharomyces uvarum*. Nature Communications 5.

12. Fay, J.C., and Benavides, J.A. (2005). Evidence for domesticated and wild populations of *Saccharomyces cerevisiae*. PLoS Genet 1, e5.

13. Gallone, B., Steensels, J., Prahl, T., Soriaga, L., Saels, V., Herrera-Malaver, B., Merlevede, A., Roncoroni, M., Voordeckers, K., Miraglia, L., et al. (2016). Domestication and divergence of *Saccharomyces cerevisiae* beer yeasts. Cell 166, 1397–1410.e16.

14. Legras, J.-L., Merdinoglu, D., Cornuet, J.-M., and Karst, F. (2007). Bread, beer and wine: *Saccharomyces cerevisiae* diversity reflects human history. Molecular Ecology 16, 2091–2102.

15. Libkind, D., Hittinger, C.T., Valério, E., Gonçalves, C., Dover, J., Johnston, M., Gonçalves, P., and Sampaio, J.P. (2011). Microbe domestication and the identification of the wild genetic stock of lager-brewing yeast. PNAS 108, 14539–14544.

16. Marsit, S., Mena, A., Bigey, F., Sauvage, F.-X., Couloux, A., Guy, J., Legras, J.-L., Barrio, E., Dequin, S., and Galeote, V. (2015). Evolutionary advantage conferred by an eukaryote-to-eukaryote gene transfer event in wine yeasts. Molecular Biology and Evolution 32, 1695–1707.

17. Novo, M., Bigey, F., Beyne, E., Galeote, V., Gavory, F., Mallet, S., Cambon, B., Legras, J.-L., Wincker, P., Casaregola, S., et al. (2009). Eukaryote-to-eukaryote gene transfer events revealed by the genome sequence of the wine yeast *Saccharomyces cerevisiae* EC1118. PNAS 106, 16333–16338.

18. Peter, J., Chiara, M.D., Friedrich, A., Yue, J.-X., Pflieger, D., Bergström, A., Sigwalt, A., Barre, B., Freel, K., Llored, A., et al. (2018). Genome evolution across 1,011 *Saccharomyces cerevisiae* isolates. Nature 556, 339–344.

19. Liti, G., Carter, D.M., Moses, A.M., Warringer, J., Parts, L., James, S.A., Davey, R.P., Roberts, I.N., Burt, A., Koufopanou, V., et al. (2009). Population genomics of domestic and wild yeasts. Nature 458, 337–341.

20. Spor, A., Kvitek, D.J., Nidelet, T., Martin, J., Legrand, J., Dillmann, C., Bourgais, A., De Vienne, D., Sherlock, G., and Sicard, D. (2014). Phenotypic and genotypic convergences are influenced by historical contingency and environment in yeast. Evolution 68, 772–790.

21. Dunn, B., Paulish, T., Stanbery, A., Piotrowski, J., Koniges, G., Kroll, E., Louis, E.J., Liti, G., Sherlock, G., and Rosenzweig, F. (2013). Recurrent rearrangement during adaptive evolution in an interspecific yeast hybrid suggests a model for rapid introgression. PLoS Genetics 9, e1003366.

22. Galagan, J.E., Calvo, S.E., Cuomo, C., Ma, L.-J., Wortman, J.R., Batzoglou, S., Lee, S.-I., Baştürkmen, M., Spevak, C.C., Clutterbuck, J., et al. (2005). Sequencing of *Aspergillus nidulans* and comparative analysis with *A. fumigatus* and *A. oryzae*. Nature 438, 1105–1115.

23. Gibbons, J.G., Salichos, L., Slot, J.C., Rinker, D.C., McGary, K.L., King, J.G., Klich, M.A., Tabb, D.L., McDonald, W.H., and Rokas, A. (2012). The evolutionary imprint of domestication on genome variation and function of the filamentous fungus *Aspergillus oryzae*. Current Biology 22, 1403–1409.

24. Machida, M., Asai, K., Sano, M., Tanaka, T., Kumagai, T., Terai, G., Kusumoto, K.-I., Arima, T., Akita, O., Kashiwagi, Y., et al. (2005). Genome sequencing and analysis of *Aspergillus oryzae*. Nature 438, 1157–1161.

25. Dumas, E., Feurtey, A., de la Vega, R.C.R., Prieur, S.L., Snirc, A., Coton, M., Thierry, A., Coton, E., Piver, M.L., Roueyre, D., et al. (2020). Independent domestication events in the blue-cheese fungus *Penicillium roqueforti*. Molecular Ecology 00, 1–22.

26. Cheeseman, K., Ropars, J., Renault, P., Dupont, J., Gouzy, J., Branca, A., Abraham, A.-L., Ceppi, M., Conseiller, E., Debuchy, R., et al. (2014). Multiple recent horizontal transfers of a large genomic region in cheese making fungi. Nat Commun 5, 2876.

27. Ropars, J., Rodríguez de la Vega, R.C., VLópez-illavicencio, M., Gouzy, J., Sallet, E., Dumas, É., Lacoste, S., Debuchy, R., Dupont, J., Branca, A., et al. (2015). Adaptive horizontal gene transfers between multiple cheese-associated Fungi. Current Biology 25, 2562–2569.

28. Gillot, G., Jany, J.-L., Coton, M., Le Floch, G., Debaets, S., Ropars, J., López-Villavicencio, M., Dupont, J., Branca, A., Giraud, T., et al. (2015). Insights into *Penicillium roqueforti* morphological and genetic diversity. PLoS ONE 10, e0129849.

29. Legras, J.-L., Galeote, V., Bigey, F., Camarasa, C., Marsit, S., Nidelet, T., Sanchez, I., Couloux, A., Guy, J., Franco-Duarte, R., et al. (2018). Adaptation of *S. cerevisiae* to fermented food environments reveals remarkable genome plasticity and the footprints of domestication. Mol Biol Evol 35, 1712–1727.

30. Pitt, J.I. (1979). The genus *Penicillium* and its teleomorphic states *Eupenicillium* and *Talaromyces*. (London, UK: Academic Press).

31. Delfosse, C. (2008). Histoires de bries (Saint-Cyr-sur-Morin: Illustria Librairie des Musées).

32. Pitt, J.I., Cruickshank, R.H., and Leistner, L. (1986). *Penicillium commune*, *P. camembertii*, the origin of white cheese moulds, and the production of cyclopiazonic acid. Food Microbiology 3, 363–371.

33. Pouriau, A.-F.A. du texte (1895). La laiterie : art de traiter le lait, de fabriquer le beurre et les principaux fromages français et étrangers (5e éd.).

34. Garnier, L., Valence, F., Pawtowski, A., Auhustsinava-Galerne, L., Frotté, N., Baroncelli, R., Deniel, F., Coton, E., and Mounier, J. (2017). Diversity of spoilage fungi associated with various French dairy products. International Journal of Food Microbiology 241, 191–197.

35. Giraud, F., Giraud, T., Aguileta, G., Fournier, E., Samson, R., Cruaud, C., Lacoste, S., Ropars, J., Tellier, A., and Dupont, J. (2010). Microsatellite loci to recognize species for the cheese starter and contaminating strains associated with cheese manufacturing. International Journal of Food Microbiology 137, 204–213.

36. Visagie, C.M., Houbraken, J., Frisvad, J.C., Hong, S.-B., Klaassen, C.H.W., Perrone, G., Seifert, K.A., Varga, J., Yaguchi, T., and Samson, R.A. (2014). Identification and nomenclature of the genus *Penicillium*. Studies in Mycology 78, 343–371.

37. Hymery, N., Vasseur, V., Coton, M., Mounier, J., Jany, J.-L., Barbier, G., and Coton, E. (2014). Filamentous Fungi and Mycotoxins in Cheese: A Review. Comprehensive Reviews in Food Science and Food Safety 13, 437–456.

38. Le Bars, J. (1979). Cyclopiazonic acid production by *Penicillium camemberti* Thom and natural occurrence of this mycotoxin in cheese. Appl Environ Microbiol 38, 1052–1055.

39. Bodinaku, I., Shaffer, J., Connors, A.B., Steenwyk, J.L., Biango-Daniels, M.N., Kastman, E.K., Rokas, A., Robbat, A., and Wolfe, B.E. (2019). Rapid Phenotypic and Metabolomic Domestication of Wild Penicillium Molds on Cheese. mBio 10, e02445–19.

40. Gibbons, J.G., and Rokas, A. (2013). The function and evolution of the *Aspergillus* genome. Trends in Microbiology 21, 14–22.

41. Dyer, P.S., and O’Gorman, C.M. (2012). Sexual development and cryptic sexuality in fungi: insights from *Aspergillus* species. FEMS Microbiol. Rev. 36, 165–192.

42. Barve, M.P., Arie, T., Salimath, S.S., Muehlbauer, F.J., and Peever, T.L. (2003). Cloning and characterization of the mating type (MAT) locus from *Ascochyta rabiei* (teleomorph: *Didymella rabiei*) and a MAT phylogeny of legume-associated *Ascochyta* spp. Fungal Genetics and Biology 39, 151–167.

43. Boisard, P. (2007). Le camembert, mythe français Odile Jacob.

44. Selker, E.U., Cambareri, E.B., Jensen, B.C., and Haack, K.R. (1987). Rearrangement of duplicated DNA in specialized cells of Neurospora. Cell 51, 741–752.

45. Hashem, K.M., He, F.J., Jenner, K.H., and MacGregor, G.A. (2014). Cross-sectional survey of salt content in cheese: a major contributor to salt intake in the UK. BMJ Open 4, e005051–e005051.

46. Kritikos, G., Banzhaf, M., Herrera-Dominguez, L., Koumoutsi, A., Wartel, M., Zietek, M., and Typas, A. (2017). A tool named Iris for versatile high-throughput phenotyping in microorganisms. Nature Microbiology 2, 17014.

47. Hymery, N., Masson, F., Barbier, G., and Coton, E. (2014). Cytotoxicity and immunotoxicity of cyclopiazonic acid on human cells. Toxicol In Vitro 28, 940–947.

48. Lund, F., Filtenborg, O., and Frisvad, J.C. (1998). *Penicillium caseifulvum,* a new species found on *Penicillium roqueforti* fermented cheeses. Journal of Food Mycology 1, 95–101.

49. Barbesgaard, P., Heldt-Hansen, H.P., and Diderichsen, B. (1992). On the safety of *Aspergillus oryzae*: a review. Appl Microbiol Biotechnol 36, 569–572.

50. Gillot, G., Jany, J.-L., Poirier, E., Maillard, M.-B., Debaets, S., Thierry, A., Coton, E., and Coton, M. (2017). Functional diversity within the *Penicillium roqueforti* species. Int. J. Food Microbiol. 241, 141–150.

51. Giraud, T., Refrégier, G., Le Gac, M., de Vienne, D.M., and Hood, M.E. (2008). Speciation in fungi. Fungal Genetics and Biology 45, 791–802.

52. Le Gac, M., and Giraud, T. (2008). Existence of a pattern of reproductive character displacement in Homobasidiomycota but not in Ascomycota. Journal of Evolutionary Biology 21, 761–772.

53. Mahler, D.L., Ingram, T., Revell, L.J., and Losos, J.B. (2013). Exceptional convergence on the macroevolutionary landscape in island lizard radiations. Science 341, 292–295.

54. Thorpe, R.S., Barlow, A., Malhotra, A., and Surget-Groba, Y. (2015). Widespread parallel population adaptation to climate variation across a radiation: implications for adaptation to climate change. Mol Ecol 24, 1019–1030.

55. Hohenlohe, P.A., Bassham, S., Etter, P.D., Stiffler, N., Johnson, E.A., and Cresko, W.A. (2010). Population Genomics of Parallel Adaptation in Threespine Stickleback using Sequenced RAD Tags. PLoS Genetics 6, e1000862.

56. Duboué, E.R., Keene, A.C., and Borowsky, R.L. (2011). Evolutionary convergence on sleep loss in cavefish populations. Current Biology 21, 671–676.

57. Martínez-Ainsworth, N.E., and Tenaillon, M.I. (2016). Superheroes and masterminds of plant domestication. Comptes Rendus Biologies 339, 268–273.

58. Gross, B.L., and Olsen, K.M. (2010). Genetic perspectives on crop domestication. Trends in Plant Science 15, 529–537.

59. Decker, M., and Nielsen, P.V. (2005). The inhibitory effect of *Penicillium camemberti* and *Geotrichum candidum* on the associated funga of white mould cheese. International Journal of Food Microbiology 104, 51–60.

60. Gouyon, P.-H., Leriche, H., Civard, A., Reeves, H., and Hulot, N. (2010). Aux origines de l’environnement (Fayard).

61. Vavilov, N.I., Vavylov, M.I., Vavílov, N.Í., and Dorofeev, V.F. (1992). Origin and Geography of Cultivated Plants (Cambridge University Press).

62. Glass, N.L., and Donaldson, G.C. (1995). Development of primer sets designed for use with the PCR to amplify conserved genes from filamentous ascomycetes. Appl Environ Microbiol 61, 1323–1330.

63. Koren, S., Walenz, B.P., Berlin, K., Miller, J.R., Bergman, N.H., and Phillippy, A.M. (2017). Canu: scalable and accurate long-read assembly via adaptive k-mer weighting and repeat separation. Genome Res. 27, 722–736.

64. Walker, B.J., Abeel, T., Shea, T., Priest, M., Abouelliel, A., Sakthikumar, S., Cuomo, C.A., Zeng, Q., Wortman, J., Young, S.K., et al. (2014). Pilon: An Integrated Tool for Comprehensive Microbial Variant Detection and Genome Assembly Improvement. PLoS ONE 9, e112963.

65. Bolger, A.M., Lohse, M., and Usadel, B. (2014). Trimmomatic: a flexible trimmer for Illumina sequence data. Bioinformatics 30, 2114–2120.

66. Langmead, B., and Salzberg, S.L. (2012). Fast gapped-read alignment with Bowtie 2. Nat. Methods 9, 357–359.

67. McKenna, A., Hanna, M., Banks, E., Sivachenko, A., Cibulskis, K., Kernytsky, A., Garimella, K., Altshuler, D., Gabriel, S., Daly, M., et al. (2010). The Genome Analysis Toolkit: a MapReduce framework for analyzing next-generation DNA sequencing data. Genome Res. 20, 1297–1303.

68. Pfeifer, B., Wittelsbürger, U., Ramos-Onsins, S.E., and Lercher, M.J. (2014). PopGenome: An Efficient Swiss Army Knife for Population Genomic Analyses in R. Mol Biol Evol 31, 1929–1936.

69. Korneliussen, T.S., Albrechtsen, A., and Nielsen, R. (2014). ANGSD: Analysis of Next Generation Sequencing Data. BMC Bioinformatics 15, 356.

70. Schliep, K.P. (2011). phangorn: phylogenetic analysis in R. Bioinformatics 27, 592–593.

71. Minh, B.Q., Schmidt, H.A., Chernomor, O., Schrempf, D., Woodhams, M.D., von Haeseler, A., and Lanfear, R. (2020). IQ-TREE 2: New Models and Efficient Methods for Phylogenetic Inference in the Genomic Era. Mol Biol Evol 37, 1530–1534.

72. Kalyaanamoorthy, S., Minh, B.Q., Wong, T.K.F., von Haeseler, A., and Jermiin, L.S. (2017). ModelFinder: fast model selection for accurate phylogenetic estimates. Nature Methods 14, 587–589.

73. Anisimova, M., Gil, M., Dufayard, J.-F., Dessimoz, C., and Gascuel, O. (2011). Survey of Branch Support Methods Demonstrates Accuracy, Power, and Robustness of Fast Likelihood-based Approximation Schemes. Syst Biol 60, 685–699.

74. Minh, B.Q., Nguyen, M.A.T., and von Haeseler, A. (2013). Ultrafast approximation for phylogenetic bootstrap. Mol. Biol. Evol. 30, 1188–1195.

75. Hill, T.W., and Kafer, E. (2001). Improved protocols for *Aspergillus* minimal medium: trace element and minimal medium salt stock solutions. Fungal Genetics Reports 48, 20–21.

76. Peterson, R.A., and Cavanaugh, J.E. (2019). Ordered quantile normalization: a semiparametric transformation built for the cross-validation era. Journal of Applied Statistics 0, 1–16.

77. Porcher, E., Giraud, T., Goldringer, I., and Lavigne, C. (2004). Experimental demonstration of a causal relationship between heterogeneity of selection and genetic differentiation in quantitative traits. Evolution 58, 1434–1445.

